# A phylogenetically informed comparative analysis of sexual testosterone dimorphism across mammals in relation to paternal care and sexual size dimorphism

**DOI:** 10.64898/2026.05.20.726499

**Authors:** B.N. Laubi, J. M. Burkart, E.P. Willems, C. P. van Schaik

## Abstract

Within species, male testosterone is often linked to mating competition and paternal care, suggesting that sex differences in endogenous testosterone values across mammals may covary with broader reproductive strategies. Using a structured literature search, we compiled 63 studies, spanning 31 non-human species and 9 human populations, reporting endogenous, non-experimentally manipulated testosterone values for both adult males and females within the same population and context. From these studies, we calculated male-to-female testosterone ratios, and analysed these data using Bayesian phylogenetic multilevel models. We tested whether testosterone dimorphism was associated with paternal care and sexual size dimorphism while accounting for sampling matrix, assay method, breeding context, and wild versus captive setting. Across non-human mammals, neither paternal care nor sexual size dimorphism (indexing competition) showed a clear association with testosterone ratios, and the same pattern emerged in the primate-only subset. By contrast, sampling matrix was consistently associated with testosterone dimorphism across all analyses, with lower male-to-female ratios in non-blood than in blood-based measures. In primates, testosterone ratios were also lower in captive than in wild populations, although this pattern was not clearly supported in the broader non-human dataset. In the human-only analysis, testosterone ratios did not clearly differ between industrialized and small-scale societies, whereas the matrix effect remained evident. Overall, our results suggest that sampling matrix is a major source of variation even for ratio-based measures, highlighting the need for caution when inferring between-species endocrine differences from studies using different substrates. More broadly, directly comparable, non-experimentally manipulated testosterone data for both sexes remain rare across mammals, limiting comparative inference.

## Introduction

Testosterone is the principal circulating androgen in most mammals and plays a central role in reproductive physiology in both sexes, including supporting the development and maintenance of primary and secondary sex characteristics and reproductive function in males, and contributing to ovarian function in females (e.g., Das et al., 2023; Walters, 2015). Beyond physiology, it has also been repeatedly linked to variation in mating, competitive and other social behaviours, and a large literature shows that testosterone levels, particularly in males, shift in response to social and reproductive challenges (Archer, 2006; Moore et al., 2020; Muller & Wrangham, 2004; Wingfield et al., 1990). Transient increases in testosterone can facilitate such behaviours, but sustained elevations are thought to carry costs for immune function, survival and parental investment, implying trade-offs between mating effort and other fitness components (Crespi et al., 2025; Hau, 2007; Muehlenbein & Bribiescas, 2005; Muller, 2017). Although both sexes produce testosterone throughout much of their lives, in most species adult males show higher circulating testosterone concentrations than adult females, consistent with stronger male-biased mating competition (Rege et al., 2019; Vernasco & Moore, 2020). At the same time, the magnitude of this sex difference varies widely within and across species, underscoring that testosterone is not simply a static “male hormone”.

One broad source of this cross-species variation may be differences in the intensity of sexual selection and mating competition. Across mammals, sexual size dimorphism is often used as an index of sex differences in mating competition, with more strongly male-biased body size differences typically occurring in systems where males compete more intensely for mates (Plavcan, 2012; Weckerly, 1998). Because testosterone is closely linked to competitive and mating-related phenotypes, species with greater sexual size dimorphism may therefore exhibit more strongly male-biased testosterone concentrations (e.g., Wingfield et al., 1990).

Another major source of variation in testosterone dimorphism may be differences in male parental investment, particularly the extent to which males contribute to infant care. In men, experimental and longitudinal studies indicate that testosterone often decreases with pair formation and active fathering, whereas competitive or extra-pair contexts are associated with elevated levels (Gettler, McDade, Agustin, et al., 2011; Gettler, McDade, Feranil, et al., 2011; Gray et al., 2002; Grebe et al., 2019; McIntyre et al., 2006; Van Der Meij et al., 2012). Comparable patterns have been documented in several nonhuman mammals, where transitions into paternal care or alloparental roles are accompanied by reductions or altered dynamics in male testosterone (Nunes et al., 2000, 2001; Wynne-Edwards, 2001; Ziegler et al., 2004, 2009).

Together, these findings highlight parenting as a context in which testosterone regulation is especially responsive to shifting reproductive effort.

Mammals differ strikingly in how male parental effort is expressed, ranging from species in which males provide little or no care to systems with intensive paternal and alloparental investment (Balshine, 2012; Rymer & Pillay, 2018). At one end of this continuum are many polygynous or solitary breeders, where males compete for mating opportunities and offspring care is largely maternal (e.g., Boness & Bowen, 1996; van Noordwijk & van Schaik, 2005). At the other end are cooperative and biparental breeders, such as callitrichid primates, some canids and rodents, and many human societies, in which adult males and other group members routinely carry, protect, provision or otherwise care for dependent young (e.g., Gettler et al., 2020; Gromov, 2020; Sparkman et al., 2011; Yamamoto, 2005). In such systems, male reproductive success depends not only on competition for mates but also on successful offspring rearing and coordination within a caregiving unit. These differences in male reproductive roles are thus expected to be associated with different endocrine profiles: more strongly male-biased testosterone regimes in systems with little male care, and more attenuated or flexible sex differences in species where males are integral carers within cooperative or biparental breeding systems. However, variation in male parental care is not independent of variation in mating competition, which makes it difficult to separate predictions based on parenting effort from those based on competition.

Empirical evidence is broadly consistent with this reasoning but remains fragmented and taxonomically patchy, with relevant data scattered across taxa and study designs. In cooperatively breeding common marmosets, for instance, we found relatively small sex differences in hair testosterone, a pattern qualitatively consistent with expectations for systems with high male caregiving effort and a mostly monogamous mating system (Laubi et al., in press). By contrast, studies of polygynous Old World primates often report pronounced male-biased testosterone, although direct quantitative comparisons across studies are complicated by methodological heterogeneity (e.g., Rege et al., 2019; Sonnweber et al., 2022). Similar between-species contrasts have been reported in rodents, including lower male testosterone in a monogamous, biparental vole than in a closely related polygynous species, consistent with broader evidence that androgen profiles covary with paternal role and social system in biparental rodents (Klein et al., 1999; Saltzman et al., 2017). Humans provide a complementary line of evidence at the population level: several studies report substantially lower average male testosterone in small-scale subsistence populations, including hunter-gatherers and forager-horticulturalists, than in industrialised populations, patterns that have been discussed in relation to high levels of alloparental support and investment in parenting (Alvergne et al., 2009; Bribiescas, 1996; Ellison, 2002; Muller et al., 2009).

Despite this conceptual and empirical work, comparative inference remains difficult because hormone data are rarely measured on a common scale across studies. Even when studies nominally use the same assay, absolute testosterone concentrations can differ due to laboratory effects, protocol differences, calibration and extraction procedures, and operator-specific variation (Calamari et al., 2020; French et al., 2019; Van Uytfanghe et al., 2005; Vesper et al., 2009). In addition, comparability depends on sampling matrix and context because different matrices (serum, plasma, saliva, urine, faeces, hair) capture different fractions or integration windows and testosterone varies systematically with time of day, breeding season, and reproductive state (Behringer & Deschner, 2017; Brambilla et al., 2009; Smith et al., 2013; Urlacher et al., 2022). As a result, absolute values are often not comparable across studies and taxa. A practical way forward is therefore to use within-study, scale-free measures of sex difference, such as male-to-female testosterone ratios, which can be derived whenever both sexes are measured in the same population and context. Cross-species tests additionally require phylogenetically informed analyses because species are not independent datapoints. Indeed, comparative work on birds found that proposed links between seasonal-maximum male-to-female testosterone ratios and life-history traits weakened or disappeared after controlling for shared ancestry (Goymann & Wingfield, 2014; Møller et al., 2005).

Building on this approach, we present a cross-taxonomic, phylogenetically informed synthesis of sex differences in testosterone across mammals, with particular emphasis on primates. Restricting the dataset to studies that sampled both sexes in the same population and context, we quantify sex differences using male-to-female testosterone ratios. We then test whether the magnitude of testosterone dimorphism covaries with male contribution to infant care and with sexual size dimorphism across mammals. Finally, we examine whether these associations are also evident within primates and whether human populations differing in socioecology and paternal/alloparental investment conform to the broader mammalian pattern.

## Methods

### Literature search

We conducted a structured literature search in the Web of Science Core Collection (Clarivate Analytics) in September 2025 to identify studies reporting testosterone concentrations for adult males and females in mammals under non-experimental conditions, with an emphasis on primates. To facilitate taxon-specific screening and reduce irrelevant hits, we ran separate queries for non-human primates, other non-human mammals, and humans. Search strings were built from core concept blocks (testosterone, sex-related descriptors, measurement/reporting terms, and biological matrices) and combined with taxon-specific filters and exclusion terms where appropriate (see Table 1 for example search terms). Matrix terms were included to improve search specificity for manual screening. Complete queries, including query-specific filters and database settings, are reported in Supplementary Table S1. Exclusion terms and database filters were used to reduce retrieval of clinical and patient studies, and records from preprint databases were excluded. For the industrialized human query, we initially restricted the Web of Science search to LC–MS/MS-related terms and blood-based matrices to keep the number of retrieved records manageable. No language restrictions were applied at the search stage. To increase completeness, we additionally screened Google Scholar using simplified combinations of the same core concepts to identify potentially eligible records not indexed in Web of Science; in this supplementary search, we did not restrict by biological matrix or assay method.

**Table 1.** Example search terms used to construct Web of Science queries. Full database queries, including Boolean operators and field tags, are provided in Supplementary Table S1.

| Search component | Example terms |
| --- | --- |
| Hormone | testosterone |
| Taxonomic filters | “primate* OR monkey* OR ape* OR lemur*” |
| Sex-related descriptors | "male and female", "both sexes", "sex differences" |
| Measurement/reporting terms | "level*", "concentration*", "baseline", "reference range*" |
| Biological matrices | "plasma", "serum", "saliva", "urine", "feces", "hair" |

### Eligibility criteria

Studies were included if they reported testosterone concentrations for both adult males and adult females within the same species and study, measured under non-experimental conditions. Eligible studies provided sex-specific quantitative data as means or medians (with measures of variability where available) or individual-level values from which summary statistics could be calculated, reported numerically in the text or tables or extractable from figures. No restrictions were applied regarding biological matrix, and measurements from blood, saliva, urine, faeces, or hair were all considered eligible. Both wild and captive non-domesticated populations were included to maximise taxonomic and ecological coverage. We included peer-reviewed articles and academic theses where full text was available in English. Studies were excluded when values did not represent comparable adult concentrations for both sexes under the same sampling context. This included experimentally manipulated animals, domesticated laboratory strains or breeds, non-adult individuals, single-sex reports, helper-only samples in cooperatively breeding species, values reported as broad androgen immunoreactivity due to substantial assay cross-reactivity, non-comparable male and female values derived from different analytical procedures, and censored estimates based only on samples above the assay detection limit.

### Screening process

Records identified through the Web of Science search and the supplementary Google Scholar search were exported into Zotero and retained in separate collections. Duplicate records were removed using Zotero’s automated duplicate detection followed by manual checking. Screening followed a two-step procedure based on the predefined eligibility criteria. First, titles and abstracts were manually screened to exclude clearly irrelevant records (e.g., studies without testosterone measurements, non-mammalian taxa, or studies that did not include both sexes). Second, full texts were obtained for all remaining records and assessed against the eligibility criteria described above. A large language model (ChatGPT, OpenAI) was used as an auxiliary tool to help locate sex-specific numerical values in full texts and figures, but all inclusion decisions and extracted values were verified manually against the original publications. Table 2 summarises the Web of Science screening process by search category and reports the number of additional eligible studies added from Google Scholar.

**Table 2.** Study selection and inclusion outcomes for the literature search. Counts are shown separately for non-human mammals, primates, and humans. Rows summarise records retrieved and screened from the Web of Science search, followed by additional studies identified through a supplementary manual search in Google Scholar and included after applying the same eligibility criteria.

| Stage | Mammals | Primates | Humans | Total |
| --- | --- | --- | --- | --- |
| Records retrieved<br>(Web of Science) | 579 | 587 | 55 | 1221 |
| Duplicates removed | 5 | 5 | 0 | 10 |
| Title/abstract screened | 574 | 582 | 55 | 1211 |
| Excluded at title/abstract | 489 | 415 | 18 | 922 |
| Full texts sought | 84 | 167 | 37 | 288 |
| Full texts not found | 22 | 16 | 1 | 39 |
| Full texts assessed | 62 | 151 | 36 | 249 |
| Excluded after full-text screening | 48 | 134 | 31 | 213 |
| **Studies included (Web of Science)** | **14** | **17** | **5** | **36** |
| **Studies added (Google Scholar)** | **7** | **5** | **4** | **16** |
| **Total studies included** | **21** | **22** | **9** | **52** |

### Data extraction

For each eligible study, data were entered into a standardized Excel spreadsheet. We recorded basic study descriptors (species, population setting, breeding context) and key methodological information (biological matrix and assay type). Population setting was coded as wild or captive for nonhuman mammals, and as industrialized or small-scale society for human studies. Breeding context was coded as breeding when samples were collected during a reported breeding season or when sampling included both breeding and non-breeding periods. It was coded as non-breeding when samples were collected outside the breeding season or when no discrete breeding season was reported for non-seasonal breeders. This variable should therefore be interpreted as a coarse descriptor of whether sampling occurred during, or included, a reported breeding season, rather than as a precise classification of individual reproductive state. Wherever available, we recorded sex-specific means and/or medians of testosterone concentrations and sample sizes; when only individual values were given, we calculated the corresponding summary statistics. When numerical data were not explicitly reported, sex-specific values were extracted from published figures using WebPlotDigitizer (version 5). Data extraction was performed manually using the predefined spreadsheet structure. The compiled study-level dataset, including extracted testosterone values, study descriptors, and derived male-to-female ratios, is provided in Supplementary Table S2.

### Comparative predictor variables

For each species in the comparative dataset, we compiled information on paternal care and body mass dimorphism. Paternal care was coded using the binary classification provided by Heldstab et al. (2019), which was derived from the comparative parental care dataset of Isler and van Schaik (2012). Species were classified according to the presence or absence of direct male infant care (i.e., carrying, provisioning), rather than protection or affiliative contact in the absence of sustained direct care (e.g., huddling). For species not covered by Heldstab et al. (2019) or Isler and van Schaik (2012), paternal care was coded using the same decision rule based on published species accounts and primary field studies. Body mass dimorphism was calculated as the ratio of average adult male to average adult female body mass. These values were primarily derived from the species-level sex-specific body mass data reported by Tombak et al. (2024); for species not included in that dataset, corresponding estimates were obtained through supplementary literature searches. Human populations were considered separately, and paternal care was coded at the population level, with populations classified as either small-scale or industrialized as a coarse proxy for socioecological context and expected paternal involvement. The resulting classifications for all species and populations are listed in Supplementary Tables S3 & S4.

### Data synthesis

For each study, we calculated the male-to-female testosterone ratio from sex-specific mean concentrations. For two human studies that reported testosterone as median with interquartile range (IQR), we converted these summaries to approximate means to harmonise effect-size calculation across sources (Kische et al., 2016; Wang et al., 2022). Specifically, we estimated the mean as (*Q*1 + median + *Q*3)/3, where *Q*1and *Q*3are the 25th and 75th percentiles, respectively. An additional human study reported sex-specific medians with 5th and 95th percentiles rather than quartiles (Schiffer et al., 2023); because these percentiles capture the distribution tails and are not directly comparable to IQR-based summaries, we used the reported medians directly to derive the male-to-female ratio in this case.

### Statistical and phylogenetic analysis

All analyses were conducted in R version 4.3.3 (R Core Team, 2024). We analysed male-to-female testosterone ratios using Bayesian phylogenetic multilevel models in the ‘brms’ package (Bürkner, 2017). Because ratios were positive and right-skewed, models were fitted with a Gamma error distribution and log link. All non-human models included sampling matrix (blood vs. nonblood), assay method (immunoassay vs. mass spectrometry), breeding context (breeding vs. non-breeding), and setting (wild vs. captive) as fixed effects, with study identity included as a random intercept to account for non-independence among estimates derived from the same source. Shared ancestry was modelled by including species as a phylogenetic random effect, using a species-level correlation matrix derived from a pruned subtree of the Open Tree of Life synthetic tree (Hinchliff et al., 2015), retrieved using ‘rotl’ (Michonneau et al., 2016). Species names were matched to the tree taxonomy prior to pruning, and the resulting phylogeny was converted to a correlation matrix for use in ‘brms’. We fitted two separate models for the non-human comparative dataset (N = 56 observations, 43 studies, 31 species) that differed in their focal biological predictor: binary paternal care and sexual size dimorphism. These predictors covaried strongly across species, with paternal-care species largely restricted to the low-dimorphism range, and were therefore analysed in separate models rather than interpreted as independent effects within a single combined model. Sexual size dimorphism was z-transformed prior to analysis to aid interpretability and improve sampling efficiency. To assess whether these patterns differed within primates, the same model set was repeated in a primate-only subset (N = 27 observations, 22 studies, 13 species). Human data were analysed separately from the non-human comparative models and did not include a phylogenetic term. Because the human subset was very small (N = 10) and assay method was strongly confounded with sampling matrix, the final human model included population category (small-scale vs. industrialized) and sampling matrix as predictors, with study identity retained as a random intercept. Human analyses could not account for population-level sexual size dimorphism because comparable estimates were not available across studies. Models were estimated with weakly informative priors and four Markov chain Monte Carlo chains of 4,000 iterations each, with 2,000 warmup iterations. Convergence was assessed using Rhat values and effective sample sizes, and model fit was evaluated using posterior predictive checks and Pareto-smoothed leave-one-out cross-validation (PSIS-LOO), implemented for ‘brms’ models via the ‘loo’ package (Vehtari et al., 2017). The distribution of observations across categorical predictors in the non-human mammal, primate-only, and human datasets is summarized in Supplementary Table S5.

## Results

We tested the predictions that paternal care would be associated with lower testosterone ratios and that sexual size dimorphism would be associated with higher ratios. In phylogenetic models of non-human animals, we found no clear evidence that either focal biological predictor explained male-to-female testosterone ratios (paternal care: beta = 0.26, 95% CrI = -0.56 to 1.11; sexual size dimorphism: beta = 0.07, 95% CrI = -0.36 to 0.49). Across both models, however, testosterone ratios were consistently lower in non-blood than blood samples (paternal care model: beta = -1.63, 95% CrI = -2.27 to -0.98; dimorphism model: beta = -1.62, 95% CrI = -2.24 to -0.96; Figure 3), whereas effects of breeding context, assay method, and setting remained uncertain.

Restricting the analysis to primates yielded a similar overall pattern. Paternal care (beta = 0.00, 95% CrI = -1.00 to 0.96) and sexual size dimorphism (beta = 0.12, 95% CrI = -0.26 to 0.55) likewise showed no clear association with male-to-female testosterone ratios. As in the broader non-human dataset, testosterone ratios were consistently lower in non-blood than blood samples across both primate models (paternal care model: beta = -2.04, 95% CrI = -2.57 to -1.46; dimorphism model: beta = -1.98, 95% CrI = -2.54 to -1.37). In addition, unlike in the broader non-human analyses, testosterone ratios were consistently lower in captive than wild primates (paternal care model: beta = -1.01, 95% CrI = -1.54 to -0.43; dimorphism model: beta = -1.01, 95% CrI = -1.53 to -0.46), whereas effects of assay method and breeding context remained uncertain.

In the human-only model, male-to-female testosterone ratios did not clearly differ between small-scale and industrialized populations (beta = 0.52, 95% CrI = -0.32 to 1.34). By contrast, testosterone ratios were again lower in non-blood than blood samples (beta = -2.43, 95% CrI = -3.25 to -1.04). Sampling matrix was therefore the only predictor that showed a consistent association with testosterone ratios across all analyses (Figure 2).

**Figure 1.**
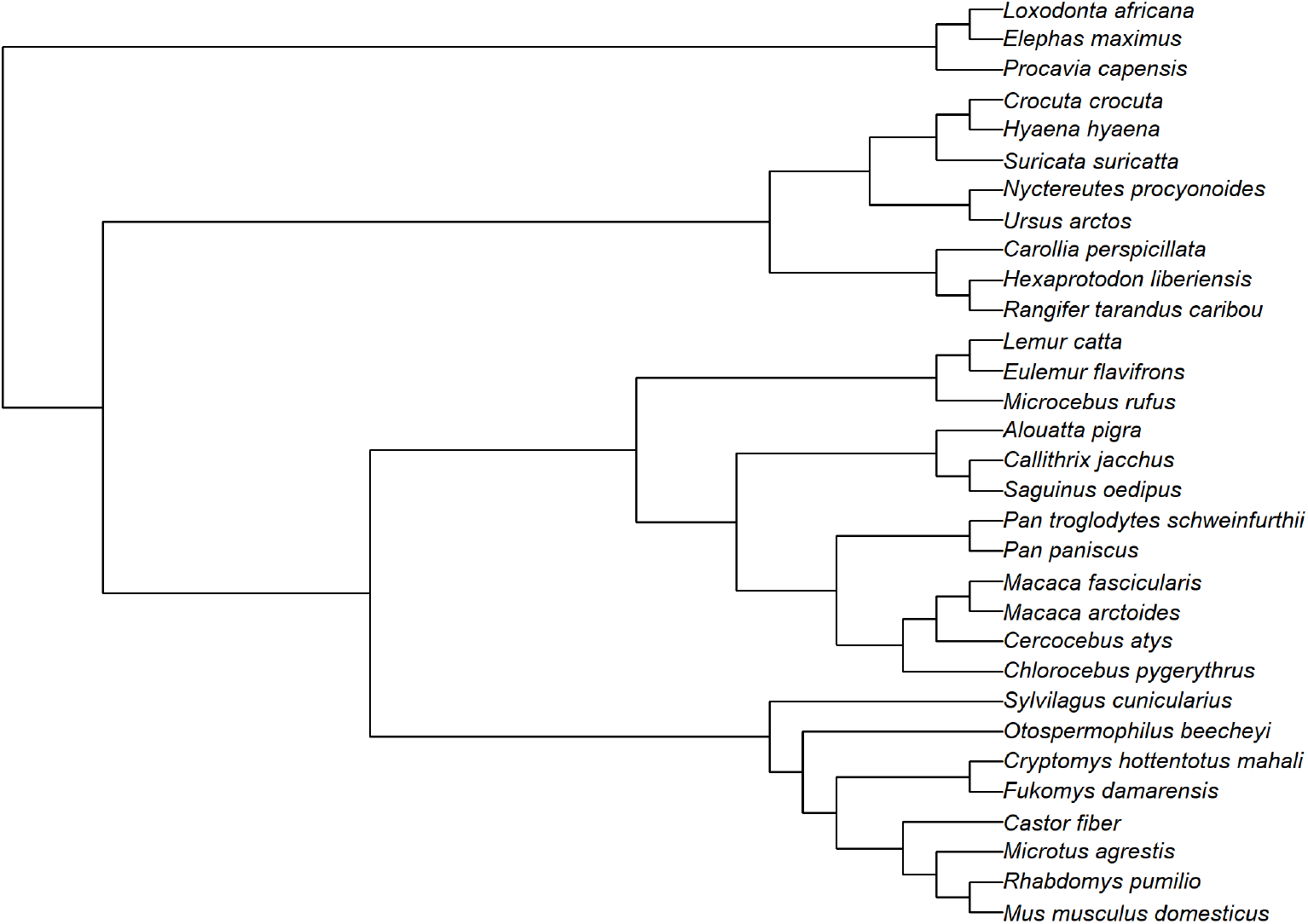
Phylogenetic tree of the non-human species (N = 31) included in the comparative analyses. The tree was derived from a pruned subtree of the Open Tree of Life synthetic tree (Hinchliff et al., 2015) and served as the basis for the phylogenetic correlation matrix used in the Bayesian comparative models.

**Figure 2.**
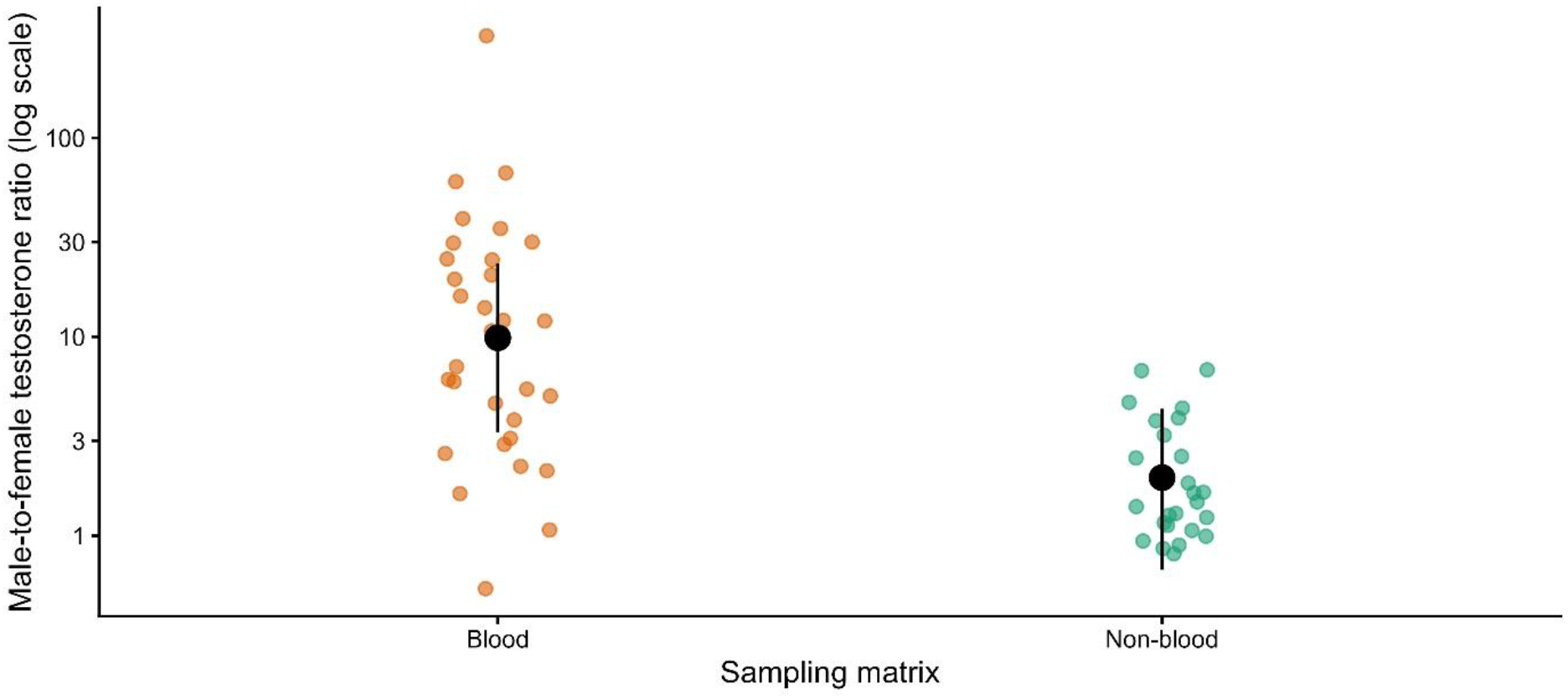
Male-to-female testosterone ratios by sampling matrix in non-human animals. Coloured points represent raw observations, and black points with 95% credible intervals represent fitted values from a reduced phylogenetic model including sampling matrix, assay method, breeding context, and setting, but excluding the focal biological predictors. The y-axis is shown on a log scale. Model estimates consistently indicated lower male-to-female testosterone ratios in non-blood than in blood-based samples across analyses and datasets.

**Figure 3.**
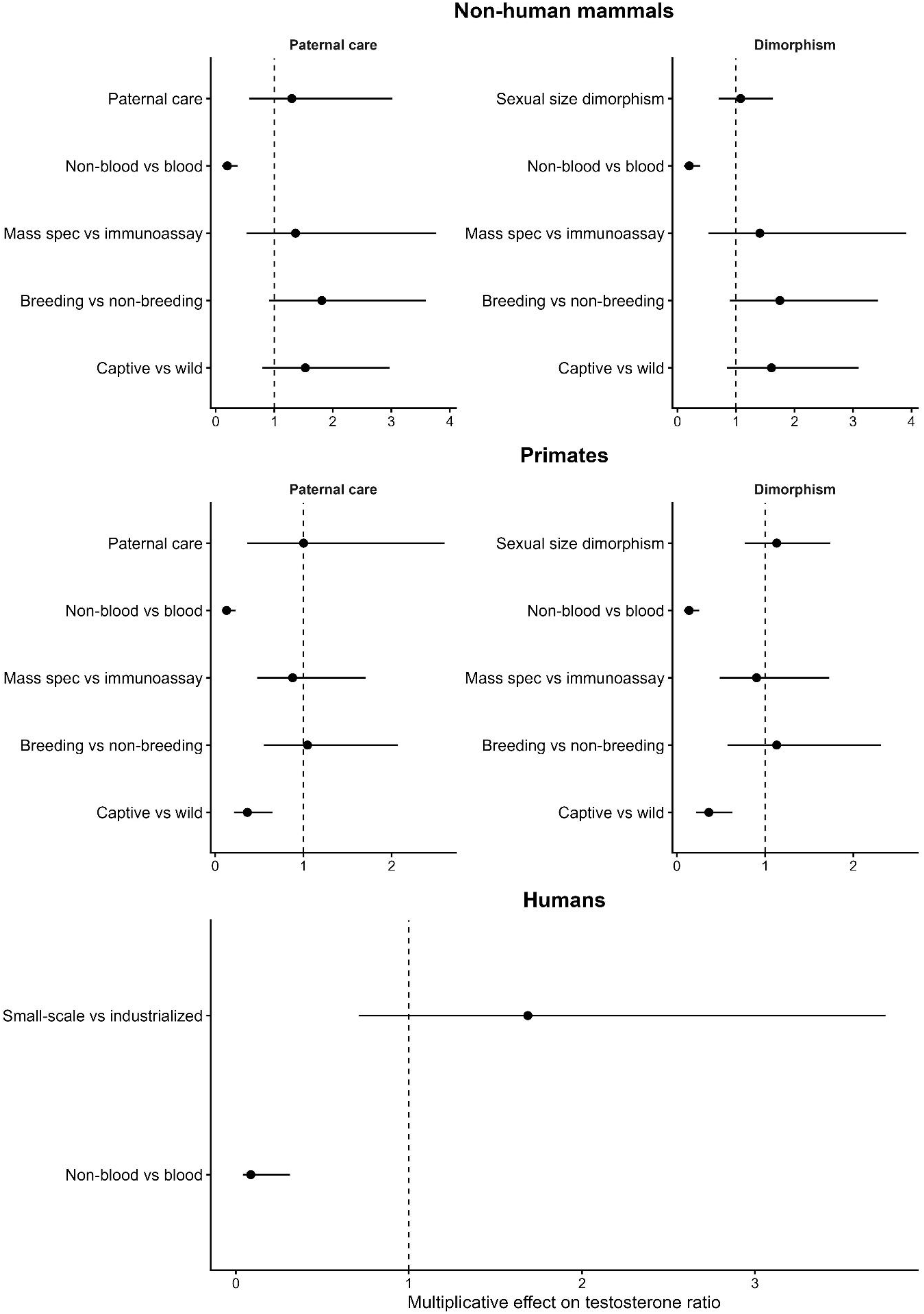
Posterior estimates and 95% credible intervals for fixed effects in the non-human mammal, primate-only, and human models. Estimates are shown on the response scale as multiplicative effects on the male-to-female testosterone ratio, with the dashed vertical line indicating no effect. Values below 1 indicate lower testosterone ratios, and values above 1 indicate higher testosterone ratios relative to the reference category.

## Discussion

We synthesized sex differences in testosterone measures across mammals using male-to-female ratios calculated from studies that reported testosterone values for both sexes in the same population and sampling context, and analysed these data with Bayesian phylogenetic multilevel models. Across non-human mammals, we found no clear evidence that either paternal care or sexual size dimorphism explained variation in testosterone ratios, and the same was true in the primate-only subset. In contrast, sampling matrix showed a consistent association across all analyses, with testosterone ratios markedly lower in non-blood than blood-based measures. In primates, we additionally observed lower testosterone ratios in captive than wild populations, whereas this effect was not clearly supported in the full non-human mammal dataset. In the human-only model, population category (small-scale vs industrialized) showed no clear association with testosterone ratios, while the matrix effect persisted, further underscoring sampling matrix as the most robust predictor of estimated sex differences in this dataset.

We predicted that species with paternal care would show less pronounced male-biased testosterone ratios, reflecting a trade-off between mating effort and investment in caregiving. Contrary to this expectation, paternal care was not clearly associated with male-to-female testosterone ratios in the non-human or primate-only analyses. This absence of a detectable effect does not necessarily contradict within-species evidence that male testosterone often decreases with pair bonding and active care (e.g., Grebe et al., 2019; Ziegler et al., 2009). Rather, it suggests that any cross-species signal is weak relative to the heterogeneity of the available data. Alternatively, it may reflect the fact that limited data availability forced us to code paternal care as a binary trait. This coarse classification may have been insufficient to capture endocrine differences linked to variable caregiving effort and social context.

We also expected sexual size dimorphism to predict male-to-female testosterone ratios, given that strongly male-biased sex differences in body size are often interpreted as reflecting more intense male mating competition (Plavcan, 2012; Weckerly, 1998). However, sexual size dimorphism showed no clear association with testosterone ratios in either dataset. One explanation, consistent with recent comparative work, is that sexual size dimorphism is an imperfect proxy for mating competition across mammals and may reflect a mixture of selective pressures, which could weaken any simple relationship with testosterone ratios (Cassini, 2020; Isaac, 2005; Jones & Sheard, 2023; Tombak et al., 2024). In addition, our ratio-based approach necessarily incorporates variation in both sexes. This is important because, although less extensively studied than in males, female testosterone may vary in ways that do not necessarily covary with male testosterone. Available evidence indicates that testosterone and other androgens can vary with reproductive state, dominance, aggression, and female-female competition (Beehner et al., 2005; Drea et al., 2021; Koren et al., 2006). Such partly independent variation in female testosterone could weaken associations between testosterone ratios and competition-related predictors such as sexual size dimorphism.

More generally, both predictors are unevenly distributed across taxa and covary with broader reproductive strategies, limiting the overlap in trait combinations available for comparative inference. Combined with substantial heterogeneity in sampling matrix, assay approach, and sampling context across studies, this may have obscured modest biological associations even when using within-study ratios.

The clearest result across all analyses was the effect of sampling matrix, with male-to-female testosterone ratios being markedly lower in non-blood than in blood-based measures. Crucially, because sampling matrix altered male-to-female ratios and not only absolute concentrations, the effect cannot be reduced to a simple conversion factor that affects males and females equally. Several non-mutually exclusive mechanisms may explain this pattern. Different matrices capture different fractions and temporal integration windows of androgen production. Blood-based measures typically quantify circulating testosterone in serum or plasma over short time scales, often reflecting total testosterone (i.e. the combined protein-bound and free hormone in circulation), whereas non-blood matrices may primarily reflect free hormone (saliva), metabolites (urine, faeces), or longer-term integrated deposition (hair) (Behringer & Deschner, 2017; Urlacher et al., 2022). This may be especially relevant if males and females differ in hormone-binding proteins, steroid metabolism, conjugation, or excretion pathways, such that the relationship between circulating testosterone and non-blood measures is not identical across sexes (Goymann, 2012; Schiffer et al., 2019). Matrix effects may also be methodological, because different matrices require different extraction procedures, validation steps, and analytical assumptions. These differences may affect estimated sex differences if assay performance is not equivalent across the concentration ranges or metabolite profiles typical of males and females. For example, cross-reactivity, incomplete recovery, calibration differences, or detection limits may bias one sex more strongly than the other, particularly in complex non-blood matrices (Vesper et al., 2009; Vesper & Botelho, 2010). A further possibility is that the ratios are affected by sex-specific temporal dynamics in testosterone. Because non-blood matrices often integrate endocrine activity over longer periods than blood, they may attenuate short-term fluctuations that are more apparent in circulating measures, potentially contributing to both lower and less variable male-to-female ratios. This interpretation is consistent with evidence from humans, where measurements in men are more strongly influenced by circadian and other short-term sources of variation, whereas variation in women is often discussed more in relation to menstrual-cycle phase (Brambilla et al., 2009; Kanakis et al., 2019). Although the present dataset cannot disentangle these mechanisms, the consistency of the matrix effect shows that substrate choice shapes inferred testosterone dimorphism and should be treated as a substantial component of comparative inference rather than a minor nuisance factor. Comparative studies should therefore model matrix explicitly and avoid treating different substrates as directly interchangeable.

In the primate-only analyses, we additionally found lower male-to-female testosterone ratios in captive than in wild populations. This pattern may indicate that captivity alters the ecological and social conditions under which testosterone is regulated. In primates, testosterone is closely tied to male mating effort and competition, and captive settings can affect both behaviour and hormone production in ways that are not straightforward to predict (Muller, 2017). Captive primates often experience restricted space, altered social housing, and husbandry-driven environments, whereas wild primates are additionally exposed to ecological and social challenges such as predation, food shortage, parasitism, intense aggressive interactions, and energetic constraints (De Oliveira Terceiro et al., 2021; Novak et al., 2013). One specific possibility is that captive management attenuates male-male competition, for example by controlling group composition or separating incompatible males, thereby reducing a key social stimulus for male testosterone. Although our models included matrix, assay method, breeding context, and setting as additive predictors, the primate-only dataset was too sparse to test whether the captive-wild difference depended on matrix, assay method, or breeding context. Biological and methodological explanations for the captive-wild difference therefore remain difficult to disentangle. Because this effect was evident in primates but not clearly supported in the full non-human mammal dataset, and because sample sizes were limited, it should be regarded as suggestive rather than conclusive.

In the human-only analysis, we found no clear evidence that population category predicted male-to-female testosterone ratios. Interpretation is limited, however, by the small number of available studies and by the necessarily coarse grouping of populations. The small-scale category combined populations with different subsistence systems, including hunter-gatherer and horticulturalist groups, whereas the industrialized category combined rural and urban settings. Moreover, human population categories capture many socioecological differences beyond paternal-care context, including variation in energetic condition, pathogen exposure, and energy expenditure that may influence testosterone levels (Trumble et al., 2023). Thus, these categories should be interpreted as coarse proxies rather than direct tests of paternal-care effects. Nevertheless, the matrix effect was again recovered in the human model, with lower testosterone ratios in non-blood than in blood-based measures, reinforcing that sampling matrix remained the most consistent predictor of estimated sex differences even in this small and heterogeneous subset.

Several limitations should be considered when interpreting these findings. First, although our literature search identified 1,221 records through Web of Science and a further 16 studies through Google Scholar, only 52 studies met the criteria for inclusion in the final dataset because few reported adult testosterone values for males and females in the same population and sampling context. This limited availability of sex-specific testosterone data highlights a broader gap in the literature and contributed to the sparse taxonomic coverage and uneven distribution of predictor combinations in our analyses. Although we accounted for phylogenetic relatedness and included study identity as a random effect, the dataset remained heterogeneous, with estimates derived from different laboratories, assay platforms, extraction procedures, and sampling protocols. Data were also sparse for some combinations of predictors, particularly breeding context and captive setting, which limited power and increased sensitivity to influential observations. Breeding context was necessarily coded coarsely because the dataset was too limited to distinguish breeding-season, non-breeding-season, non-seasonal and mixed sampling contexts. This coding may have diluted the contrast between reproductive contexts and obscured biologically meaningful seasonal variation in testosterone ratios. Thus, breeding context should be interpreted as a broad control variable rather than as a precise test of mating, rearing, or non-reproductive periods. Finally, our binary coding of paternal care necessarily collapsed important variation in the form and intensity of male care, while the sexual size dimorphism estimates were compiled from multiple sources and may not always capture within-species variation.

Overall, our synthesis indicates that sampling matrix is the most robust predictor of estimated male-to-female testosterone ratios across the available mammalian literature, whereas paternal care and sexual size dimorphism were not clearly associated with testosterone dimorphism at this comparative scale. This does not necessarily mean that these biological factors are irrelevant, but rather that their effects are difficult to detect against the background of sparse taxonomic coverage, uneven trait distributions, and substantial methodological heterogeneity. Future work would benefit from more standardized comparative datasets using matched sampling contexts, consistent matrices, and ideally analytically comparable methods such as LC-MS/MS, as well as from more informative biological predictors such as continuous measures of male care. More broadly, this study underscores how much comparative insight still depends on a basic prerequisite that remains surprisingly rare in the literature, namely directly comparable testosterone data for both sexes.

## Acknowledgements

We are grateful to Maria Granell Ruiz for generously sharing unpublished testosterone data for vervet monkeys included in this comparative analysis.

## References

Alvergne, A., Faurie, C., & Raymond, M. (2009). Variation in testosterone levels and male reproductive effort: Insight from a polygynous human population. Hormones and Behavior, 56(5), 491–497. 10.1016/j.yhbeh.2009.07.013

Archer, J. (2006). Testosterone and human aggression: An evaluation of the challenge hypothesis. Neuroscience & Biobehavioral Reviews, 30(3), 319–345. 10.1016/j.neubiorev.2004.12.007

Balshine, S. (2012). Patterns of parental care in vertebrates. In M. Kölliker, The Evolution of Parental Care (pp. 62–80). Oxford University Press. 10.1093/acprof:oso/9780199692576.003.0004

Beehner, J. C., Phillips-Conroy, J. E., & Whitten, P. L. (2005). Female testosterone, dominance rank, and aggression in an Ethiopian population of hybrid baboons. American Journal of Primatology, 67(1), 101–119. 10.1002/ajp.20172

Behringer, V., & Deschner, T. (2017). Non-invasive monitoring of physiological markers in primates. Hormones and Behavior, 91, 3–18. 10.1016/j.yhbeh.2017.02.001

Boness, D. J., & Bowen, W. D. (1996). The Evolution of Maternal Care in Pinnipeds. BioScience, 46(9), 645–654. 10.2307/1312894

Brambilla, D. J., Matsumoto, A. M., Araujo, A. B., & McKinlay, J. B. (2009). The Effect of Diurnal Variation on Clinical Measurement of Serum Testosterone and Other Sex Hormone Levels in Men. The Journal of Clinical Endocrinology & Metabolism, 94(3), 907–913. 10.1210/jc.2008-1902

Bribiescas, R. G. (1996). Testosterone levels among Aché hunter-gatherer men: A functional interpretation of population variation among adult males. Human Nature, 7(2), 163–188.

Bürkner, P.-C. (2017). brms: An R Package for Bayesian Multilevel Models Using Stan. Journal of Statistical Software, 80(1). 10.18637/jss.v080.i01

Calamari, C. V., Viau, P., Nichi, M., Martins, G. S., Sobral, G., Mangueira Dias, J. H., & Alvarenga De Oliveira, C. (2020). Hair as an alternative noninvasive matrix: Sources of variation in testosterone levels. Domestic Animal Endocrinology, 72, 106477. 10.1016/j.domaniend.2020.106477

Cassini, M. H. (2020). A mixed model of the evolution of polygyny and sexual size dimorphism in mammals. Mammal Review, 50(1), 112–120. 10.1111/mam.12171

Crespi, B. J., Bushell, A., & Dinsdale, N. (2025). Testosterone mediates life-history trade-offs in female mammals. Biological Reviews, 100(2), 871–891. 10.1111/brv.13166

Das, P. K., Sejian, V., Mukherjee, J., & Banerjee, D. (Eds). (2023). Functional morphology of the male reproductive system. In Textbook of Veterinary Physiology (pp. 441–476). Springer Nature Singapore. 10.1007/978-981-19-9410-4

De Oliveira Terceiro, F. E., Arruda, M. D. F., Van Schaik, C. P., Araújo, A., & Burkart, J. M. (2021). Higher social tolerance in wild versus captive common marmosets: The role of interdependence. Scientific Reports, 11(1), 825. 10.1038/s41598-020-80632-3

Drea, C. M., Davies, C. S., Greene, L. K., Mitchell, J., Blondel, D. V., Shearer, C. L., Feldblum, J. T., Dimac-Stohl, K. A., Smyth-Kabay, K. N., & Clutton-Brock, T. H. (2021). An intergenerational androgenic mechanism of female intrasexual competition in the cooperatively breeding meerkat. Nature Communications, 12(1), 7332. 10.1038/s41467-021-27496-x

Ellison, P. T. (2002). Population variation in age-related decline in male salivary testosterone. Human Reproduction, 17(12), 3251–3253. 10.1093/humrep/17.12.3251

French, D., Drees, J., Stone, J. A., Holmes, D. T., & Van Der Gugten, J. G. (2019). Comparison of four clinically validated testosterone LC-MS/MS assays: Harmonization is an attainable goal. Clinical Mass Spectrometry, 11, 12–20. 10.1016/j.clinms.2018.11.005

Gettler, L. T., Boyette, A. H., & Rosenbaum, S. (2020). Broadening Perspectives on the Evolution of Human Paternal Care and Fathers’ Effects on Children. Annual Review of Anthropology, 49(1), 141–160. 10.1146/annurev-anthro-102218-011216

Gettler, L. T., McDade, T. W., Agustin, S. S., & Kuzawa, C. W. (2011). Short-term changes in fathers’ hormones during father–child play: Impacts of paternal attitudes and experience. Hormones and Behavior, 60(5), 599–606. 10.1016/j.yhbeh.2011.08.009

Gettler, L. T., McDade, T. W., Feranil, A. B., & Kuzawa, C. W. (2011). Longitudinal evidence that fatherhood decreases testosterone in human males. Proceedings of the National Academy of Sciences, 108(39), 16194–16199. 10.1073/pnas.1105403108

Goymann, W. (2012). On the use of non-invasive hormone research in uncontrolled, natural environments: The problem with sex, diet, metabolic rate and the individual. Methods in Ecology and Evolution, 3(4), 757–765. 10.1111/j.2041-210X.2012.00203.x

Goymann, W., & Wingfield, J. C. (2014). Male-to-female testosterone ratios, dimorphism, and life history—What does it really tell us? Behavioral Ecology, 25(4), 685–699. 10.1093/beheco/aru019

Gray, P. B., Kahlenberg, S. M., Barrett, E. S., Lipson, S. F., & Ellison, P. T. (2002). Marriage and fatherhood are associated with lower testosterone in males. Evolution and Human Behavior, 23(3), 193–201. 10.1016/S1090-5138(01)00101-5

Grebe, N. M., Sarafin, R. E., Strenth, C. R., & Zilioli, S. (2019). Pair-bonding, fatherhood, and the role of testosterone: A meta-analytic review. Neuroscience & Biobehavioral Reviews, 98, 221–233. 10.1016/j.neubiorev.2019.01.010

Gromov, V. S. (2020). Paternal care in rodents: Ultimate causation and proximate mechanisms. Russian Journal of Theriology, 19(1), 1–20. 10.15298/rusjtheriol.19.1.01

Hau, M. (2007). Regulation of male traits by testosterone: Implications for the evolution of vertebrate life histories. BioEssays, 29(2), 133–144. 10.1002/bies.20524

Heldstab, S. A., Isler, K., Burkart, J. M., & Van Schaik, C. P. (2019). Allomaternal care, brains and fertility in mammals: Who cares matters. Behavioral Ecology and Sociobiology, 73(6), 71. 10.1007/s00265-019-2684-x

Hinchliff, C. E., Smith, S. A., Allman, J. F., Burleigh, J. G., Chaudhary, R., Coghill, L. M., Crandall, K. A., Deng, J., Drew, B. T., Gazis, R., Gude, K., Hibbett, D. S., Katz, L. A., Laughinghouse, H. D., McTavish, E. J., Midford, P. E., Owen, C. L., Ree, R. H., Rees, J. A., … Cranston, K. A. (2015). Synthesis of phylogeny and taxonomy into a comprehensive tree of life. Proceedings of the National Academy of Sciences, 112(41), 12764–12769. 10.1073/pnas.1423041112

Isaac, J. L. (2005). Potential causes and life-history consequences of sexual size dimorphism in mammals. Mammal Review, 35(1), 101–115. 10.1111/j.1365-2907.2005.00045.x

Isler, K., & Van Schaik, C. P. (2012). Allomaternal care, life history and brain size evolution in mammals. Journal of Human Evolution, 63(1), 52–63. 10.1016/j.jhevol.2012.03.009

Jones, M. E., & Sheard, C. (2023). The macroevolutionary dynamics of mammalian sexual size dimorphism. Proceedings of the Royal Society B: Biological Sciences, 290(2011).

Kanakis, G. A., Tsametis, C. P., & Goulis, D. G. (2019). Measuring testosterone in women and men. Maturitas, 125, 41–44. 10.1016/j.maturitas.2019.04.203

Kische, H., Gross, S., Wallaschofski, H., Voelzke, H., Doerr, M., Nauck, M., Felix, S. B., & Haring, R. (2016). Serum androgen concentrations and subclinical measures of cardiovascular disease in men and women. ATHEROSCLEROSIS, 247, 193–200. (WOS:000372718900026). 10.1016/j.atherosclerosis.2016.02.020

Klein, S. L., Gamble, H. R., & Nelson, R. J. (1999). Role of steroid hormones in Trichinella spiralis infection among voles. American Journal of Physiology-Regulatory, Integrative and Comparative Physiology, 277(5), R1362–R1367. 10.1152/ajpregu.1999.277.5.R1362

Koren, L., Mokady, O., & Geffen, E. (2006). Elevated testosterone levels and social ranks in female rock hyrax. Hormones and Behavior, 49(4), 470–477. 10.1016/j.yhbeh.2005.10.004

Laubi, B., Glauser, G., Willems, E. P., Van Schaik, C., & Burkart, J. (in press). Hair steroid signatures of cooperative breeding in male and female common marmosets (Callithrix jacchus). International Journal of Primatology.

McIntyre, M., Gangestad, S. W., Gray, P. B., Chapman, J. F., Burnham, T. C., O’Rourke, M. T., & Thornhill, R. (2006). Romantic involvement often reduces men’s testosterone levels--but not always: The moderating role of extrapair sexual interest. Journal of Personality and Social Psychology, 91(4), 642–651. 10.1037/0022-3514.91.4.642

Michonneau, F., Brown, J. W., & Winter, D. J. (2016). rotl: An R package to interact with the Open Tree of Life data. Methods in Ecology and Evolution, 7(12), 1476–1481. 10.1111/2041-210X.12593

Møller, A. P., Garamszegi, L. Z., Gil, D., Hurtrez-Boussès, S., & Eens, M. (2005). Correlated evolution of male and female testosterone profiles in birds and its consequences. Behavioral Ecology and Sociobiology, 58(6), 534–544. 10.1007/s00265-005-0962-2

Moore, I. T., Hernandez, J., & Goymann, W. (2020). Who rises to the challenge? Testing the Challenge Hypothesis in fish, amphibians, reptiles, and mammals. Hormones and Behavior, 123, 104537. 10.1016/j.yhbeh.2019.06.001

Muehlenbein, M. P., & Bribiescas, R. G. (2005). Testosterone-mediated immune functions and male life histories. American Journal of Human Biology, 17(5), 527–558. 10.1002/ajhb.20419

Muller, M. N. (2017). Testosterone and reproductive effort in male primates. Hormones and Behavior, 91, 36–51. 10.1016/j.yhbeh.2016.09.001

Muller, M. N., Marlowe, F. W., Bugumba, R., & Ellison, P. T. (2009). Testosterone and paternal care in East African foragers and pastoralists. Proceedings of the Royal Society B: Biological Sciences, 276(1655), 347–354. 10.1098/rspb.2008.1028

Muller, M. N., & Wrangham, R. W. (2004). Dominance, aggression and testosterone in wild chimpanzees: A test of the ‘challenge hypothesis’. Animal Behaviour, 67(1), 113–123. 10.1016/j.anbehav.2003.03.013

Novak, M. A., Hamel, A. F., Kelly, B. J., Dettmer, A. M., & Meyer, J. S. (2013). Stress, the HPA axis, and nonhuman primate well-being: A review. Applied Animal Behaviour Science, 143(2–4), 135–149. 10.1016/j.applanim.2012.10.012

Nunes, S., Fite, J. E., & French, J. A. (2000). Variation in steroid hormones associated with infant care behaviour and experience in male marmosets (Callithrix kuhlii). Animal Behaviour, 60(6), 857–865. 10.1006/anbe.2000.1524

Nunes, S., Fite, J. E., Patera, K. J., & French, J. A. (2001). Interactions among Paternal Behavior, Steroid Hormones, and Parental Experience in Male Marmosets (Callithrix kuhlii). Hormones and Behavior, 39(1), 70–82. 10.1006/hbeh.2000.1631

Plavcan, J. M. (2012). Sexual Size Dimorphism, Canine Dimorphism, and Male-Male Competition in Primates: Where Do Humans Fit In? Human Nature, 23(1), 45–67. 10.1007/s12110-012-9130-3

R Core Team. (2024). R: A Language and Environment for Statistical Computing (Version 4.3.3) [Computer software]. R Foundation for Statistical Computing. https://www.R-project.org/

Rege, J., Garber, S., Conley, A. J., Elsey, R. M., Turcu, A. F., Auchus, R. J., & Rainey, W. E. (2019). Circulating 11-oxygenated androgens across species. The Journal of Steroid Biochemistry and Molecular Biology, 190, 242–249. 10.1016/j.jsbmb.2019.04.005

Rymer, T. L., & Pillay, N. (2018). An integrated understanding of paternal care in mammals: Lessons from the rodents. Journal of Zoology, 306(2), 69–76. 10.1111/jzo.12575

Saltzman, W., Harris, B. N., De Jong, T. R., Perea-Rodriguez, J. P., Horrell, N. D., Zhao, M., & Andrew, J. R. (2017). Paternal Care in Biparental Rodents: Intra- and Inter-individual Variation. Integrative and Comparative Biology, 57(3), 589–602. 10.1093/icb/icx047

Schiffer, L., Barnard, L., Baranowski, E. S., Gilligan, L. C., Taylor, A. E., Arlt, W., Shackleton, C. H. L., & Storbeck, K.-H. (2019). Human steroid biosynthesis, metabolism and excretion are differentially reflected by serum and urine steroid metabolomes: A comprehensive review. The Journal of Steroid Biochemistry and Molecular Biology, 194, 105439. 10.1016/j.jsbmb.2019.105439

Schiffer, L., Kempegowda, P., Sitch, A. J., Adaway, J. E., Shaheen, F., Ebbehoj, A., Singh, S., McTaggart, M. P., O’Reilly, M. W., Prete, A., Hawley, J. M., Keevil, B. G., Bancos, I., Taylor, A. E., & Arlt, W. (2023). Classic and 11-oxygenated androgens in serum and saliva across adulthood: A cross-sectional study analyzing the impact of age, body mass index, and diurnal and menstrual cycle variation. EUROPEAN JOURNAL OF ENDOCRINOLOGY, 188(1). (WOS:000984865500005). 10.1093/ejendo/lvac017

Smith, R. P., Coward, R. M., Kovac, J. R., & Lipshultz, L. I. (2013). The evidence for seasonal variations of testosterone in men. Maturitas, 74(3), 208–212. 10.1016/j.maturitas.2012.12.003

Sonnweber, R., Stevens, J. M. G., Hohmann, G., Deschner, T., & Behringer, V. (2022). Plasma Testosterone and Androstenedione Levels Follow the Same Sex-Specific Patterns in the Two Pan Species. Biology, 11(9), 1275. 10.3390/biology11091275

Sparkman, A. M., Adams, J., Beyer, A., Steury, T. D., Waits, L., & Murray, D. L. (2011). Helper effects on pup lifetime fitness in the cooperatively breeding red wolf (Canis rufus). Proceedings of the Royal Society B: Biological Sciences, 278(1710), 1381–1389. 10.1098/rspb.2010.1921

Tombak, K. J., Hex, S. B. S. W., & Rubenstein, D. I. (2024). New estimates indicate that males are not larger than females in most mammal species. Nature Communications, 15(1), 1872. 10.1038/s41467-024-45739-5

Trumble, B. C., Pontzer, H., Stieglitz, J., Cummings, D. K., Wood, B., Emery Thompson, M., Raichlen, D., Beheim, B., Yetish, G., Kaplan, H., & Gurven, M. (2023). Energetic costs of testosterone in two subsistence populations. American Journal of Human Biology, 35(11), e23949. 10.1002/ajhb.23949

Urlacher, S. S., Kim, E. Y., Luan, T., Young, L. J., & Adjetey, B. (2022). Minimally invasive biomarkers in human and non-human primate evolutionary biology: Tools for understanding variation and adaptation. American Journal of Human Biology, 34(11), e23811. 10.1002/ajhb.23811

Van Der Meij, L., Almela, M., Hidalgo, V., Villada, C., IJzerman, H., Van Lange, P. A. M., & Salvador, A. (2012). Testosterone and Cortisol Release among Spanish Soccer Fans Watching the 2010 World Cup Final. PLoS ONE, 7(4), e34814. 10.1371/journal.pone.0034814

van Noordwijk, M. A., & van Schaik, C. P. (2005). Development of ecological competence in Sumatran orangutans. American Journal of Physical Anthropology, 127(1), 79–94. 10.1002/ajpa.10426

Van Uytfanghe, K., Stöckl, D., Kaufman, J. M., Fiers, T., De Leenheer, A., & Thienpont, L. M. (2005). Validation of 5 routine assays for serum free testosterone with a candidate reference measurement procedure based on ultrafiltration and isotope dilution–gas chromatography–mass spectrometry. Clinical Biochemistry, 38(3), 253–261. 10.1016/j.clinbiochem.2004.12.001

Vehtari, A., Gelman, A., & Gabry, J. (2017). Practical Bayesian model evaluation using leave-one-out cross-validation and WAIC. Statistics and Computing, 27(5), 1413–1432. 10.1007/s11222-016-9696-4

Vernasco, B. J., & Moore, I. T. (2020). Testosterone as a mediator of the tradeoff between cooperation and competition in the context of cooperative reproductive behaviors. General and Comparative Endocrinology, 288, 113369. 10.1016/j.ygcen.2019.113369

Vesper, H. W., Bhasin, S., Wang, C., Tai, S. S., Dodge, L. A., Singh, R. J., Nelson, J., Ohorodnik, S., Clarke, N. J., Salameh, W. A., Parker, C. R., Razdan, R., Monsell, E. A., & Myers, G. L. (2009). Interlaboratory comparison study of serum total testoserone measurements performed by mass spectrometry methods. Steroids, 74(6), 498–503. 10.1016/j.steroids.2009.01.004

Vesper, H. W., & Botelho, J. C. (2010). Standardization of testosterone measurements in humans. JOURNAL OF STEROID BIOCHEMISTRY AND MOLECULAR BIOLOGY, 121(3–5), 513–519. (WOS:000281174000006). 10.1016/j.jsbmb.2010.03.032

Walters, K. A. (2015). Role of androgens in normal and pathological ovarian function. Reproduction, 149(4), R193–R218. 10.1530/REP-14-0517

Wang, L., Chen, G., Hou, J., Wei, D., Liu, P., Nie, L., Fan, K., Wang, J., Xu, Q., Song, Y., Wang, M., Huo, W., Jing, T., Li, W., Guo, Y., Wang, C., & Mao, Z. (2022). Ambient ozone exposure combined with residential greenness in relation to serum sex hormone levels in Chinese rural adults. ENVIRONMENTAL RESEARCH, 210. (WOS:000777212100003). 10.1016/j.envres.2022.112845

Weckerly, F. W. (1998). Sexual-Size Dimorphism: Influence of Mass and Mating Systems in the Most Dimorphic Mammals. Journal of Mammalogy, 79(1), 33–52. 10.2307/1382840

Wingfield, J. C., Hegner, R. E., Dufty, A. M., & Ball, G. F. (1990). The ‘Challenge Hypothesis’: Theoretical Implications for Patterns of Testosterone Secretion, Mating Systems, and Breeding Strategies. The American Naturalist, 136(6), 829–846.

Wynne-Edwards, K. E. (2001). Hormonal Changes in Mammalian Fathers. Hormones and Behavior, 40(2), 139–145. 10.1006/hbeh.2001.1699

Yamamoto, M. E. (2005). Infant Care in Callitrichids: Cooperation and Competition. Annual Review of Biomedical Sciences, (7), 149–160.

Ziegler, T. E., Prudom, S. L., Zahed, S. R., Parlow, A. F., & Wegner, F. (2009). Prolactin’s mediative role in male parenting in parentally experienced marmosets (Callithrix jacchus). Hormones and Behavior, 56(4), 436–443. 10.1016/j.yhbeh.2009.07.012

Ziegler, T. E., Washabaugh, K. F., & Snowdon, C. T. (2004). Responsiveness of expectant male cotton-top tamarins, Saguinus oedipus, to mate’s pregnancy. Hormones and Behavior, 45(2), 84–92. 10.1016/j.yhbeh.2003.09.003

