## Supplementary Materials for "A phylogenetically informed comparative analysis of sexual testosterone dimorphism across mammals in relation to paternal care and sexual size dimorphism"

### Supplementary Table S1. Full Web of Science Core Collection search queries (Topic field, TS)

**Date searched:** September 2025
**Database exclusions applied to all searches:** Preprint Citation Index (Exclude); Research Commons (Exclude)

#### S1a. Non-human mammals

#### Topic query (TS):

TS=(testosterone)
AND TS=(("male AND female") OR "both sex*" OR "sex difference*")
AND TS=(mammal* OR rodent* OR carnivore* OR ungulate* OR bat* OR chiroptera OR cetacean* OR pinniped* OR canid* OR felid* OR mustelid* OR bovine OR bovidae OR cervid* OR suid* OR equid* OR marsupial* OR monotreme*)
NOT TS=(primate* OR monkey* OR ape* OR lemur* OR human OR patient OR clinical OR men OR women)
AND TS=(level* OR concentration* OR measur* OR baseline OR "reference interval*" OR "reference range*")
AND TS=(adult OR adults)
AND TS=(serum OR plasma OR blood OR saliva OR urine OR feces OR faeces OR hair)

---

#### S1b. Non-human primates

#### Topic query (TS):

TS=(testosterone)
AND TS=(male AND female)
AND TS=(primate* OR monkey* OR ape* OR lemur*)
AND TS=(level* OR concentration* OR measur*)
AND NOT TS=(human OR patient OR clinical OR men OR women)

---

#### S1c. Humans (industrialized; LC–MS/MS)

#### Topic query (TS):

TS=(testosterone)
AND TS=("LC-MS/MS" OR "liquid chromatography-tandem mass spectrometry" OR "isotope dilution")
AND TS=(serum OR plasma OR blood)
AND TS=(human OR humans)
AND TS=(("men AND women") OR ("male AND female"))

---

#### S1d. Humans (small-scale)

#### Topic query (TS):

TS=(testosterone)
AND TS=(("men AND women") OR ("male AND female") OR "both sex*" OR "sex difference*")
AND TS=(forager* OR "hunter-gatherer*" OR pastoralist* OR "small-scale" OR subsistence OR indigenous OR tribal OR "traditional society")
AND TS=(adult OR adults)
AND TS=(serum OR plasma OR blood OR saliva OR saliv* OR urine OR urinary OR "dried blood" OR DBS)

**Supplementary Table S2.** Study-level metadata for observations included in the comparative analyses of baseline testosterone dimorphism.
For each observation, the table reports vertebrate group, species binomial name, common name, setting, breeding context, biological matrix, analytical method, and the reference from which testosterone values were obtained. Extracted testosterone values, sample sizes and calculated ratios are not included in the preprint supplement and will be made available upon journal publication.

| Vert_Group | Species_binom | Common_name | Setting | Context | Matrix | Method | Ref |
|---|---|---|---|---|---|---|---|
| Human | Homo sapiens | human | bayaka | nonbreeding | nonblood | immunoassay | (Gettler et al., 2023) |
| Human | Homo sapiens | human | canada | nonbreeding | nonblood | immunoassay | (Kheloui et al., 2021) |
| Human | Homo sapiens | human | germany | nonbreeding | blood | mass spec | (Kische et al., 2016) |
| Human | Homo sapiens | human | germany | nonbreeding | blood | mass spec | (Kunz et al., 2023) |
| Human | Homo sapiens | human | uk | nonbreeding | blood | mass spec | (Schiffer et al., 2023) |
| Human | Homo sapiens | human | tsimane | nonbreeding | nonblood | immunoassay | (Trumble et al., 2023) |
| Human | Homo sapiens | human | south afrika | nonbreeding | blood | immunoassay | (van der Walt et al., 1977) |
| Human | Homo sapiens | human | san (!kung) | nonbreeding | blood | immunoassay | (van der Walt et al., 1977) |
| Human | Homo sapiens | human | austria | nonbreeding | blood | mass spec | (Vierhapper et al., 1997) |
| Human | Homo sapiens | human | china | nonbreeding | blood | mass spec | (Wang et al., 2022) |
| Mammal | Carollia perspicillata | short-tailed fruit bat | wild | nonbreeding | blood | immunoassay | (Greiner et al., 2010) |
| Mammal | Castor fiber | european beaver | wild | nonbreeding | nonblood | immunoassay | (Chojnowska et al., 2015) |
| Mammal | Castor fiber | european beaver | wild | nonbreeding | nonblood | immunoassay | (Chojnowska et al., 2015) |
| Mammal | Castor fiber | european beaver | wild | nonbreeding | nonblood | immunoassay | (Chojnowska et al., 2015) |
| Mammal | Choeropsis liberiensis | pygmy hippopotamus | captive | nonbreeding | nonblood | immunoassay | (Flacke et al., 2023) |
| Mammal | Crocuta crocuta | spotted hyena | captive | nonbreeding | blood | immunoassay | (Glickman et al., 1992) |
| Mammal | Crocuta crocuta | spotted hyena | wild | nonbreeding | blood | immunoassay | (Racey & Skinner, 1979) |
| Mammal | Crocuta crocuta | spotted hyena | wild | nonbreeding | blood | immunoassay | (Vanjaarsveld & Skinner, 1991) |
| Mammal | Cryptomys hottentotus mahali | mahali mole-rat | wild | nonbreeding | blood | immunoassay | (Hart et al., 2022) |
| Mammal | Elephas maximus | asian elephant | captive | nonbreeding | blood | immunoassay | (Rasmussen et al., 1984) |
| Mammal | Elephas maximus | asian elephant | captive | breeding | blood | immunoassay | (Rasmussen et al., 1984) |
| Mammal | Fukomys damarensis | damaraland mole-rats | captive | nonbreeding | nonblood | immunoassay | (Wallace et al., 2023) |

| | | | | | | | |
|---|---|---|---|---|---|---|---|
| Mammal | Hyaena hyaena | striped hyena | wild | nonbreeding | blood | immunoassay | (Racey & Skinner, 1979) |
| Mammal | Loxodonta africana | african elephant | wild | breeding | blood | immunoassay | (Rasmussen et al., 1984) |
| Mammal | Microtus agrestis | field vole | captive | breeding | blood | immunoassay | (Helle et al., 2008) |
| Mammal | Mus musculus domesticus | house mouse | wild | breeding | nonblood | mass spec | (Carlitz et al., 2022) |
| Mammal | Nyctereutes procyonoides | raccoon dog | captive | nonbreeding | blood | immunoassay | (Fentener van Vlissingen et al., 1988) |
| Mammal | Procavia capensis | rock hyrax | wild | nonbreeding | blood | immunoassay | (Koren et al., 2006) |
| Mammal | Rangifer tarandus caribou | woodland caribou | wild | breeding | nonblood | immunoassay | (Flasko et al., 2017) |
| Mammal | Rangifer tarandus caribou | woodland caribou | wild | nonbreeding | nonblood | immunoassay | (Flasko et al., 2017) |
| Mammal | Rhabdomys pumilio | striped mouse | wild | nonbreeding | blood | immunoassay | (Schradin, 2008) |
| Mammal | Rhabdomys pumilio | striped mouse | wild | breeding | blood | immunoassay | (Schradin, 2008) |
| Mammal | Spermophilus beecheyi | california ground squirrel | wild | breeding | blood | immunoassay | (Holekamp & Talamantes, 1991) |
| Mammal | Spermophilus beecheyi | california ground squirrel | wild | nonbreeding | blood | immunoassay | (Holekamp & Talamantes, 1991) |
| Mammal | Suricata suricatta | meerkats | wild | nonbreeding | blood | immunoassay | (Davies et al., 2016) |
| Mammal | Suricata suricatta | meerkats | captive | nonbreeding | blood | immunoassay | (Swift-Gallant et al., 2015) |
| Mammal | Sylvilagus cunicularius | mexican cottontail | wild | breeding | blood | immunoassay | (Aguilar et al., 2014) |
| Mammal | Ursus arctos | brown bear | wild | nonbreeding | nonblood | immunoassay | (Cattet et al., 2018) |
| Mammal | Ursus arctos | brown bear | wild | nonbreeding | nonblood | immunoassay | (Cattet et al., 2018) |
| Primate | Alouatta pigra | black howler monkey | wild | nonbreeding | nonblood | immunoassay | (Rangel-Negrin et al., 2014) |
| Primate | Callithrix jacchus | common marmoset | captive | nonbreeding | nonblood | mass spec | (Laubi et al., in press) |
| Primate | Callithrix jacchus | common marmoset | captive | nonbreeding | nonblood | immunoassay | (Melber, 2018) |
| Primate | Cercocebus atys | sooty mangabey | captive | nonbreeding | blood | immunoassay | (Mann et al., 1983) |
| Primate | Chlorocebus pygerythrus | vervet monkey | wild | breeding | nonblood | mass spec | (Granell-Ruiz, unpublished data) |
| Primate | Chlorocebus pygerythrus | vervet monkey | wild | nonbreeding | nonblood | mass spec | (Granell-Ruiz, unpublished data) |
| Primate | Eulemur flavifrons | blue-eyed black lemurs | captive | breeding | blood | immunoassay | (Grebe et al., 2022) |
| Primate | Lemur catta | ringtailed lemur | wild | breeding | blood | immunoassay | (Drea, 2007) |

| | | | | | | | |
|---|---|---|---|---|---|---|---|
| Primate | Lemur catta | ringtailed lemur | wild | breeding | blood | immunoassay | (Drea, 2011) |
| Primate | Lemur catta | ringtailed lemur | wild | nonbreeding | nonblood | immunoassay | (Tennenhouse et al., 2017) |
| Primate | Lemur catta | ringtailed lemur | wild | nonbreeding | nonblood | immunoassay | (Von Engelhard et al., 2000) |
| Primate | Lemur catta | ringtailed lemur | wild | breeding | nonblood | immunoassay | (Von Engelhard et al., 2000) |
| Primate | Macaca arctoides | stumptail macaque | captive | nonbreeding | blood | immunoassay | (Kling & Dunne, 1976) |
| Primate | Macaca arctoides | stumptail macaque | captive | nonbreeding | blood | immunoassay | (Rhodes et al., 1994) |
| Primate | Macaca fascicularis | long-tailed macaque | captive | nonbreeding | blood | immunoassay | (Cronin & Koritnik, 1983) |
| Primate | Macaca fascicularis | long-tailed macaque | captive | nonbreeding | blood | immunoassay | (Koritnik & Marschke, 1986) |
| Primate | Macaca fascicularis | long-tailed macaque | wild | nonbreeding | blood | immunoassay | (Malaivijitnond et al., 2007) |
| Primate | Macaca fascicularis | long-tailed macaque | wild | nonbreeding | blood | immunoassay | (Malaivijitnond et al., 2007) |
| Primate | Microcebus rufus | brown mouse lemur | wild | breeding | nonblood | immunoassay | (Zohdy et al., 2014) |
| Primate | Pan paniscus | bonobo | wild | nonbreeding | nonblood | immunoassay | (Dittami et al., 2008) |
| Primate | Pan paniscus | bonobo | captive | nonbreeding | nonblood | immunoassay | (Sannen et al., 2003) |
| Primate | Pan paniscus | bonobo | captive | nonbreeding | nonblood | immunoassay | (Sannen et al., 2004) |
| Primate | Pan troglodytes schweinfurthii | chimpanzee | wild | nonbreeding | nonblood | mass spec | (Hauser et al., 2011) |
| Primate | Pan troglodytes schweinfurthii | chimpanzee | wild | nonbreeding | blood | mass spec | (Hauser et al., 2011) |
| Primate | Pan troglodytes schweinfurthii | chimpanzee | wild | nonbreeding | blood | mass spec | (Preis et al., 2011) |
| Primate | Pan troglodytes schweinfurthii | chimpanzee | wild | nonbreeding | nonblood | immunoassay | (Sannen et al., 2003) |
| Primate | Saguinus oedipus | cotton-top tamarin | captive | nonbreeding | nonblood | immunoassay | (Fontani et al., 2014) |

**Supplementary Table S3.** Species-level predictor variables used in the comparative analyses of non-human mammals and primates. For each species, the table reports vertebrate group (**Vert. Group**), genus (**Genus**), species (**Species**), common name (**Common_name**), binary paternal care classifications from **Paternal care (Isler)**, **Paternal care (Heldstab)**, and **Paternal care (Laubi)**, and body mass dimorphism ratios from **Dimorphism (Tombak)** and **Dimorphism (Laubi)**. Values in the **Laubi** columns were assigned or compiled by B.L. for species not covered by the corresponding published datasets, using the same decision rules described in the Methods. The final column (**References**) lists the sources for dimorphism estimates reported in **Dimorphism (Laubi)** that were not obtained from Tombak et al. (2024)

| Vert. Group | Genus | Species | Common_name | Paternal care (Isler) | Paternal care (Heldstab) | Paternal care (Laubi) | Dimorphism (Tombak) | Dimorphism (Laubi) | References |
|---|---|---|---|---|---|---|---|---|---|
| Mammal | Carollia | perspicillata | short-tailed fruit bat | NA | NA | 0 | 1.008 | NA | |
| Mammal | Castor | fiber | european beaver | NA | NA | 1 | NA | 0.910 | (Graf et al., 2018) |
| Mammal | Choeropsis | liberiensis | pygmy hippopotamus | NA | NA | 0 | NA | 1.045 | (Meireles et al., 2025) |
| Mammal | Crocuta | crocuta | spotted hyena | 0 | 0 | 0 | NA | 0.881 | Animal Diversity Web. *Crocuta crocuta*. Accessed 20 May 2026. |
| Mammal | Cryptomys | hottentotus mahali | mahali mole-rat | NA | NA | 1 | 1.048 | NA | |
| Mammal | Elephas | maximus | asian elephant | 0 | 0 | 0 | NA | 1.333 | IELC / San Diego Zoo Wildlife Alliance Library Guides. Asian elephant. Accessed 20 May 2026. |
| Mammal | Fukomys | damarensis | damaraland mole-rats | NA | NA | 1 | NA | 1.104 | (Šumbera et al., 2026) |
| Mammal | Hyaena | hyaena | striped hyena | 0 | 0 | 1 | NA | 1.133 | (Houssaye, 2025) |
| Mammal | Loxodonta | africana | african elephant | 0 | 0 | 0 | NA | 1.970 | (Laws, 1966) |
| Mammal | Microtus | agrestis | field vole | NA | NA | 0 | 1.221 | NA | |
| Mammal | Mus | musculus domesticus | house mouse | NA | NA | 0 | 1.032 | NA | |
| Mammal | Nyctereutes | procyonoides | raccoon dog | 0 | 1 | 1 | NA | 1.136 | (Mulder, 2012) |
| Mammal | Procavia | capensis | rock hyrax | 0 | 0 | 0 | NA | 1.111 | Animal Diversity Web. *Procavia capensis*. Accessed 20 May 2026. |
| Mammal | Rangifer | tarandus caribou | woodland caribou | 0 | 0 | 0 | 1.222 | NA | |
| Mammal | Rhabdomys | pumilio | striped mouse | 0 | 0 | 0 | 1.213 | NA | |

| | | | | | | | | | |
|---|---|---|---|---|---|---|---|---|---|
| Mammal | Spermophilus | beecheyi | california ground squirrel | 0 | 0 | 0 | NA | 1.176 | Animal Diversity Web. *Spermophilus beecheyi*. Accessed 20 May 2026. |
| Mammal | Suricata | suricatta | meerkats | NA | 1 | 1 | NA | 1.015 | Animal Diversity Web. *Suricata sucicatta*. Accessed 20 May 2026. |
| Mammal | Sylvilagus | cunicularius | mexican cottontail | NA | NA | 0 | NA | 0.827 | (Lorenzo et al., 2026) |
| Mammal | Ursus | arctos | brown bear | 0 | 0 | 0 | NA | 1.879 | Animal Diversity Web. *Ursus arctos*. Accessed 20 May 2026. |
| Primate | Alouatta | pigra | black howler monkey | NA | NA | 0 | NA | 1.338 | (Kelaita et al., 2011) |
| Primate | Callithrix | jacchus | common marmoset | 1 | 1 | 1 | 0.973 | NA | |
| Primate | Cercocebus | atys | sooty mangabey | 0 | 0 | 0 | NA | 1.855 | Primate Info Net. Sooty mangabey. Accessed 20 May 2026. |
| Primate | Chlorocebus | pygerythrus | vervet monkey | NA | NA | 0 | NA | 1.341 | Primate Info Net. Vervet monkey. Accessed 20 May 2026. |
| Primate | Eulemur | flavifrons | blue-eyed black lemurs | NA | NA | 0 | 1.014 | NA | |
| Primate | Lemur | catta | ringtailed lemur | 0 | 0 | 0 | 1.01 | NA | |
| Primate | Macaca | fascicularis | long-tailed macaque | 0 | 0 | 0 | NA | 1.500 | (Fooden, 2006) |
| Primate | Macaca | arctoides | stumptail macaque | 0 | 0 | 0 | NA | 1.566 | (Larson, 1978) |
| Primate | Microcebus | rufus | brown mouse lemur | NA | 0 | 0 | NA | 0.916 | (Randrianambinina et al., 2003) |
| Primate | Pan | paniscus | bonobo | 0 | 1 | 0 | NA | 1.245 | (Zihlman & Bolter, 2015) |
| Primate | Pan | troglodytes schweinfurthii | chimpanzee | 0 | 1 | 0 | 1.225 | NA | |
| Primate | Saguinus | oedipus | cotton-top tamarin | 1 | 1 | 1 | 0.953 | NA | |

**Supplementary Table S4.** Population-level information used in the human comparative analyses. For each population, the table reports provenance and population type, as used to classify populations in the human-only models.

| Vert. Group | Genus | Species | Common_name | Provenance | Population type |
|---|---|---|---|---|---|
| Human | Homo | sapiens | human | austria | industrialized |
| Human | Homo | sapiens | human | bayaka | small-scale |
| Human | Homo | sapiens | human | canada | industrialized |
| Human | Homo | sapiens | human | china | industrialized |
| Human | Homo | sapiens | human | germany | industrialized |
| Human | Homo | sapiens | human | san (!kung) | small-scale |
| Human | Homo | sapiens | human | south afrika | industrialized |
| Human | Homo | sapiens | human | tsimane | small-scale |
| Human | Homo | sapiens | human | uk | industrialized |

**Supplementary Table S5.** Distribution of study-level observations across categorical predictors in the analytical datasets. For the non-human mammal, primate-only, and human subsets, the table reports the number of observations by sampling matrix, analytical method, breeding context, setting, paternal care classification, and population type, where applicable

| Dataset | Predictor | Category | N |
|---|---|---|---|
| Non-human mammals | Matrix | Blood | 31 |
| Non-human mammals | Matrix | Non-blood | 25 |
| Non-human mammals | Method | Immunoassay | 49 |
| Non-human mammals | Method | Mass spectrometry | 7 |
| Non-human mammals | Breeding context | Non-breeding | 42 |
| Non-human mammals | Breeding context | Breeding | 14 |
| Non-human mammals | Setting | Wild | 37 |
| Non-human mammals | Setting | Captive | 19 |
| Non-human mammals | Paternal care | Absent | 44 |
| Non-human mammals | Paternal care | Present | 12 |
| Primates | Matrix | Blood | 12 |
| Primates | Matrix | Non-blood | 15 |
| Primates | Method | Immunoassay | 21 |
| Primates | Method | Mass spectrometry | 6 |
| Primates | Breeding context | Non-breeding | 21 |
| Primates | Breeding context | Breeding | 6 |
| Primates | Setting | Wild | 16 |
| Primates | Setting | Captive | 11 |
| Primates | Paternal care | Absent | 24 |
| Primates | Paternal care | Present | 3 |
| Humans | Matrix | Blood | 7 |
| Humans | Matrix | Non-blood | 3 |
| Humans | Population type | Industrialized | 7 |
| Humans | Population type | Small-scale | 3 |

**Supplementary References**


Aguilar, F., Roedel, H. G., Vazquez, J., Nicolas, L., Rodriguez-Martinez, L., Bautista, A., & Martinez-Gomez, M. (2014). Seasonal changes in testosterone levels in wild Mexican cottontails Sylvilagus cunicularius. *MAMMALIAN BIOLOGY*, *79*(4), 225–229. (WOS:000338410700001). https://doi.org/10.1016/j.mambio.2014.02.002

Carlitz, E. H. D., Lindholm, A. K., Gao, W., Kirschbaum, C., & Konig, B. (2022). Steroid hormones in hair and fresh wounds reveal sex specific costs of reproductive engagement and reproductive success in wild house mice (Mus musculus domesticus). *HORMONES AND BEHAVIOR*, *138*. (WOS:000758028000001). https://doi.org/10.1016/j.yhbeh.2021.105102

Cattet, M., Stenhouse, G. B., Boulanger, J., Janz, D. M., Kapronczai, L., Swenson, J. E., & Zedrosser, A. (2018). Can concentrations of steroid hormones in brown bear hair reveal age class? *CONSERVATION PHYSIOLOGY*, *6*. (WOS:000423734400001). https://doi.org/10.1093/conphys/coy001

Chojnowska, K., Czerwinska, J., Kaminski, T., Kaminska, B., Panasiewicz, G., Kurzynska, A., & Bogacka, I. (2015). Sex- and seasonally related changes in plasma gonadotropins and sex steroids concentration in the European beaver (Castor fiber). *European Journal of Wildlife Research*, *61*(6), 807–811. https://doi.org/10.1007/s10344-015-0955-z

Cronin, M. J., & Koritnik, D. R. (1983). Dopamine receptors of the monkey anterior pituitary in various endocrine states. *Endocrinology*, *112*(2), 618–623. (WOS:A1983PZ00300029). https://doi.org/10.1210/endo-112-2-618

Davies, C. S., Smyth, K. N., Greene, L. K., Walsh, D. A., Mitchell, J., Clutton-Brock, T., & Drea, C. M. (2016). Exceptional endocrine profiles characterise the meerkat: Sex, status, and reproductive patterns. *Scientific Reports*, *6*(1), 35492. https://doi.org/10.1038/srep35492

Dittami, J., Katina, S., Moestl, E., Eriksson, J., Machatschke, I. H., & Hohmann, G. (2008). Urinary androgens and cortisol metabolites in field-sampled bonobos (Pan paniscus). *GENERAL AND COMPARATIVE ENDOCRINOLOGY*, *155*(3), 552–557. (WOS:000253272800009). https://doi.org/10.1016/j.ygcen.2007.08.009

Drea, C. M. (2007). Sex and seasonal differences in aggression and steroid secretion in Lemur catta: Are socially dominant females hormonally 'masculinized'? *Hormones and Behavior*, *51*(4), 555–567. https://doi.org/10.1016/j.yhbeh.2007.02.006

Drea, C. M. (2011). Endocrine correlates of pregnancy in the ring-tailed lemur (Lemur catta): Implications for the masculinization of daughters. *HORMONES AND BEHAVIOR*, *59*(4), 417–427. (WOS:000290086100001). https://doi.org/10.1016/j.yhbeh.2010.09.011

Fentener van Vlissingen, J. M., Blankenstein, M. A., Thijssen, J. H. H., Colenbrander, B., Verbruggen, A. J. E. P., & Wensing, C. J. G. (1988). Familial male pseudohermaphroditism and testicular descent in the raccoon dog *Nytereutes*. *Anatomical Record*, *222*(4), 350–356. (BCI:BCI198987065056).

Flacke, G. L., Penfold, L. M., Schwarzenberger, F., Martin, G. B., Rosales-Nieto, C. A., & Paris, M. C. J. (2023). Non-invasive assessment of fecal glucocorticoid and androgen metabolites in the pygmy hippopotamus (Choeropsis liberiensis). *GENERAL AND COMPARATIVE ENDOCRINOLOGY*, *341*. (WOS:001035761400001). https://doi.org/10.1016/j.ygcen.2023.114338

Flasko, A., Manseau, M., Mastromonaco, G., Bradley, M., Neufeld, L., & Wilson, P. (2017). Fecal DNA, hormones, and pellet morphometrics as a noninvasive method to estimate age class: An application to wild populations of Central Mountain and Boreal woodland caribou (Rangifer tarandus caribou). *CANADIAN JOURNAL OF ZOOLOGY*, *95*(5), 311–321. (WOS:000400979400002). https://doi.org/10.1139/cjz-2016-0070

Fontani, S., Vaglio, S., Beghelli, V., Mattioli, M., Bacci, S., & Accorsi, P. A. (2014). Fecal Concentrations of Cortisol, Testosterone, and Progesterone in Cotton-Top Tamarins Housed in Different Zoological Parks: Relationships Among Physiological Data, Environmental Conditions, and Behavioral Patterns. *Journal of Applied Animal Welfare Science*, *17*(3), 228–252. https://doi.org/10.1080/10888705.2014.916173

Fooden, J. (2006). Comparative Review of Fascicularis-group Species of Macaques (primates: Macaca). *Fieldiana Zoology*, *107*. https://doi.org/10.3158/0015-0754(2006)107%5B1:CROFSM%5D2.0.CO;2

Gettler, L. T., Samson, D. R., Kilius, E., Sarma, M. S., Miegakanda, V., Lew-Levy, S., & Boyette, A. H. (2023). Hormone physiology and sleep dynamics among BaYaka foragers of the Congo Basin: Gendered associations between nighttime activity, testosterone, and cortisol. *Hormones and Behavior*, *155*, 105422. https://doi.org/10.1016/j.yhbeh.2023.105422

Glickman, S., Frank, L., Pavgi, S., & Licht, P. (1992). HORMONAL CORRELATES OF MASCULINIZATION IN FEMALE SPOTTED HYAENAS (CROCUTA-CROCUTA) .1. INFANCY TO SEXUAL MATURITY. *JOURNAL OF REPRODUCTION AND FERTILITY*, *95*(2), 451–462. (WOS:A1992JF07900013).

Graf, P. M., Wilson, R. P., Sanchez, L. C., Hackländer, K., & Rosell, F. (2018). Diving behavior in a free-living, semi-aquatic herbivore, the Eurasian beaver *Castor fiber*. *Ecology and Evolution*, *8*(2), 997–1008. https://doi.org/10.1002/ece3.3726

Grebe, N. M., Sheikh, A., & Drea, C. M. (2022). Integrating the female masculinization and challenge hypotheses: Female dominance, male deference, and seasonal hormone fluctuations in adult blue-eyed black lemurs (Eulemur flavifrons)*. *HORMONES AND BEHAVIOR*, *139*. (WOS:000758043400001). https://doi.org/10.1016/j.yhbeh.2022.105108

Greiner, S., Stefanski, V., Dehnhard, M., & Voigt, C. C. (2010). Plasma testosterone levels decrease after activation of skin immune system in a free-ranging mammal. *General and Comparative Endocrinology*, *168*(3), 466–473. https://doi.org/10.1016/j.ygcen.2010.06.008

Hart, D. W., Van Vuuren, A. K. J., Erasmus, A., Süess, T., Hagenah, N., Ganswindt, A., & Bennett, N. C. (2022). The endocrine control of reproductive suppression in an aseasonally breeding social subterranean rodent, the Mahali mole-rat (Cryptomys hottentotus mahali). *Hormones and Behavior*, *142*, 105155. https://doi.org/10.1016/j.yhbeh.2022.105155

Hauser, B., Mugisha, L., Preis, A., & Deschner, T. (2011). LC–MS analysis of androgen metabolites in serum and urine from east African chimpanzees (Pan troglodytes schweinfurthii). *General and Comparative Endocrinology*, *170*(1), 92–98. https://doi.org/10.1016/j.ygcen.2010.09.012

Helle, S., Laaksonen, T., Adamsson, A., Paranko, J., & Huitu, O. (2008). Female field voles with high testosterone and glucose levels produce male-biased litters. *Animal Behaviour*, *75*(3), 1031–1039. https://doi.org/10.1016/j.anbehav.2007.08.015

Holekamp, K. E., & Talamantes, F. (1991). Seasonal-variation in circulating testosterone and estrogens of wild-caught California ground-squirrels (<i>Spermophilus beecheyi<i/>). *Journal of Reproduction and Fertility*, *93*(2), 415–425. (WOS:A1991HA96600020).

Houssaye, F. (2025). *Best Practise Guidelines: Striped Hyena*. EAZA. https://doi.org/10.61024/BPGStripedhyenaEN

Kelaita, M., Dias, P. A. D., Aguilar-Cucurachi, Ma. D. S., Canales-Espinosa, D., & Cortés-Ortiz, L. (2011). Impact of intrasexual selection on sexual dimorphism and testes size in the Mexican howler monkeys *Alouatta palliata* and *A. pigra*. *American Journal of Physical Anthropology*, *146*(2), 179–187. https://doi.org/10.1002/ajpa.21559

Kheloui, S., Brouillard, A., Rossi, M., Marin, M.-F., Mendrek, A., Paquette, D., & Juster, R.-P. (2021). Exploring the sex and gender correlates of cognitive sex differences. *ACTA PSYCHOLOGICA*, *221*. (WOS:000722001300010). https://doi.org/10.1016/j.actpsy.2021.103452

Kische, H., Gross, S., Wallaschofski, H., Voelzke, H., Doerr, M., Nauck, M., Felix, S. B., & Haring, R. (2016). Serum androgen concentrations and subclinical measures of cardiovascular disease in men and women. *ATHEROSCLEROSIS*, *247*, 193–200. (WOS:000372718900026). https://doi.org/10.1016/j.atherosclerosis.2016.02.020

Kling, A., & Dunne, K. (1976). Social-environmental factors affecting behavior and plasma testosterone in normal and amygdala lesioned *M.-speciosa*. *Primates*, *17*(1), 23–42. (BCI:BCI197662000208). https://doi.org/10.1007/BF02381564

Koren, L., Mokady, O., & Geffen, E. (2006). Elevated testosterone levels and social ranks in female rock hyrax. *Hormones and Behavior*, *49*(4), 470–477. https://doi.org/10.1016/j.yhbeh.2005.10.004

Koritnik, D. R., & Marschke, K. B. (1986). Sex steroid hormone-binding globulin levels and free 17-beta-estradiol and testosterone in cynomolgus monkeys during different reproductive states. *Journal of Steroid Biochemistry and Molecular Biology*, *25*(1), 135–141. (WOS:A1986D679200019). https://doi.org/10.1016/0022-4731(86)90292-X

Kunz, S., Wang, X., Ferrari, U., Drey, M., Theodoropoulou, M., Schilbach, K., Reincke, M., Heier, M., Peters, A., Koenig, W., Zeller, T., Thorand, B., & Bidlingmaier, M. (2023). Age- and sex-adjusted reference intervals for steroid hormones measured by liquid chromatography–tandem mass spectrometry using a widely available kit. *Endocrine Connections*, *13*(1), e230225. https://doi.org/10.1530/EC-23-0225

Larson, S. G. (1978). Scaling of organ weights in *Macaca arctoides*. *American Journal of Physical Anthropology*, *49*(1), 95–102. https://doi.org/10.1002/ajpa.1330490115

Laubi, B., Glauser, G., Willems, E. P., Van Schaik, C., & Burkart, J. (in press). Hair steroid signatures of cooperative breeding in male and female common marmosets (*Callithrix jacchus*). *International Journal of Primatology*.

Laws, R. M. (1966). Age criteria for the African elephant, Loxodonta a. Africana. *African Journal of Ecology*, *4*(1), 1–37. https://doi.org/https://doi.org/10.1111/j.1365-2028.1966.tb00878.x

Lorenzo, C., Vázquez, J., & Rodríguez-Martínez, L. (2026). Mexican Cottontail Sylvilagus cunicularius (Waterhouse, 1848). In C. Lorenzo Monterrubio & J. M. Mora (Eds), *Mammals of Middle and South America: Lagomorpha* (pp. 1–16). Springer Nature Switzerland. https://doi.org/10.1007/978-3-031-31564-0_6-1

Malaivijitnond, S., Hamada, Y., Suryobroto, B., & Takenaka, O. (2007). Female long-tailed macaques with scrotum-like structure. *AMERICAN JOURNAL OF PRIMATOLOGY*, *69*(7), 721–735. (WOS:000247669400001). https://doi.org/10.1002/ajp.20380

Mann, D. R., Castracane, V. D., McLaughlin, F., Gould, K. G., & Collins, D. C. (1983). Developmental patterns of serum luteinizing hormone, gonadal and adrenal steroids in the sooty mangabey (*Cercocebus atys*). *Biology of Reproduction*, *28*(2), 279–284. (WOS:A1983QE77400004). https://doi.org/10.1095/biolreprod28.2.279

Meireles, J., Hahn-Klimroth, M., Müller, D., Dierkes, P., & Clauss, M. (2025). Body mass of adult zoo hippos Hippopotamidae and how they compare to data from populations in the wild. *Journal of Zoo and Aquarium Research*, *13*(2), 108–116. https://doi.org/10.19227/jzar.v13i2.905

Melber, T. N. (2018). *Testosterone and Behavior in Female Marmosets*. (PQDT:60922932).

Mulder, J. L. (2012). A review of the ecology of the raccoon dog (Nyctereutes procyonoides) in Europe. *Lutra Interior*, *55*(2), 101–127.

Preis, A., Mugisha, L., Hauser, B., Weltring, A., & Deschner, T. (2011). Androgen and androgen metabolite levels in serum and urine of East African chimpanzees (Pan troglodytes schweinfurthii): Comparison of EIA and LC-MS analyses. *GENERAL AND COMPARATIVE ENDOCRINOLOGY*, *174*(3), 335–343. (WOS:000297536100012). https://doi.org/10.1016/j.ygcen.2011.09.010

Racey, P. A., & Skinner, J. D. (1979). Endocrine aspects of sexual mimicry in spotted hyaenas *Cocuta crocuta*. *Journal of Zoology*, *187*(MAR), 315–326. (WOS:A1979GM79800004).

Randrianambinina, B., Rakotondravony, D., Radespiel, U., & Zimmermann, E. (2003). Seasonal changes in general activity, body mass and reproduction of two small nocturnal primates: A comparison of the golden brown mouse lemur ( Microcebus ravelobensis ) in Northwestern Madagascar and the brown mouse lemur ( Microcebus rufus ) in Eastern Madagascar. *Primates*, *44*(4), 321–331. https://doi.org/10.1007/s10329-003-0046-8

Rangel-Negrin, A., Flores-Escobar, E., Chavira, R., Canales-Espinosa, D., & Dias, P. A. D. (2014). Physiological and analytical validations of fecal steroid hormone measures in black howler monkeys. *PRIMATES*, *55*(4), 459–465. (WOS:000343142200001). https://doi.org/10.1007/s10329-014-0432-4

Rasmussen, L. E., Buss, I. O., Hess, D. L., & Schmidt, M. J. (1984). Testosterone and Dihydrotestosterone Concentrations in Elephant Serum and Temporal Gland Secretions. *Biology of Reproduction*, *30*(2), 352–362. https://doi.org/10.1095/biolreprod30.2.352

Rhodes, L., Harper, J., Uno, H., Gaito, G., Audette-Arruda, J., Kurata, S., Berman, C., Primka, R., & Pikounis, B. (1994). The effects of finasteride (Proscar) on hair growth, hair cycle stage, and serum testosterone and dihydrotestosterone in adult male and female stumptail macaques (Macaca arctoides). *The Journal of Clinical Endocrinology & Metabolism*, *79*(4), 991–996. https://doi.org/10.1210/jcem.79.4.7962310.

Sannen, A., Heistermann, M., van Elsacker, L., Möhle, U., & Eens, M. (2003). Urinary testosterone metabolite levels on bonobos: A comparison with chimpanzees in relation to social system. *BEHAVIOUR*, *140*, 683–696. (WOS:000184773000007). https://doi.org/10.1163/156853903322149504

Sannen, A., Van Elsacker, L., Heistermann, M., & Eens, M. (2004). Urinary testosterone-metabolite levels and dominance rank in male and female bonobos (Pan paniscus). *PRIMATES*, *45*(2), 89–96. (WOS:000221014400002). https://doi.org/10.1007/s10329-003-0066-4

Schiffer, L., Kempegowda, P., Sitch, A. J., Adaway, J. E., Shaheen, F., Ebbehoj, A., Singh, S., McTaggart, M. P., O'Reilly, M. W., Prete, A., Hawley, J. M., Keevil, B. G., Bancos, I., Taylor, A. E., & Arlt, W. (2023). Classic and 11-oxygenated androgens in serum and saliva across adulthood: A cross-sectional study analyzing the impact of age, body mass index, and diurnal and menstrual cycle variation. *EUROPEAN JOURNAL OF ENDOCRINOLOGY*, *188*(1). (WOS:000984865500005). https://doi.org/10.1093/ejendo/lvac017

Schradin, C. (2008). Seasonal changes in testosterone and corticosterone levels in four social classes of a desert dwelling sociable rodent. *Hormones and Behavior*, *53*(4), 573–579. https://doi.org/10.1016/j.yhbeh.2008.01.003

Šumbera, R., Uhrová, M., Bennett, N. C., Eiseb, S. J., Faulkes, C. G., Finn, K. T., Lövy, M., Phiri, K., Van Daele, P. A. A. G., Zíková, B., & Mikula, O. (2026). Intraspecific differentiation and phylogeography of the Damaraland mole-rat Fukomys damarensis reveals rapid colonization of arid savannahs during the late Pleistocene. *Mammalian Biology*, *106*(2), 381–395. https://doi.org/10.1007/s42991-025-00546-3

Swift-Gallant, A., Mo, K., Peragine, D. E., Monks, D. A., & Holmes, M. M. (2015). Removal of reproductive suppression reveals latent sex differences in brain steroid hormone receptors in naked mole-rats, Heterocephalus glaber. *Biology of Sex Differences*, *6*(1), 31. https://doi.org/10.1186/s13293-015-0050-x

Tennenhouse, E. M., Putman, S., Boisseau, N. P., & Brown, J. L. (2017). Relationships between steroid hormones in hair and social behaviour in ring-tailed lemurs (Lemur catta). *Primates*, *58*(1), 199–209. https://doi.org/10.1007/s10329-016-0566-7

Trumble, B. C., Pontzer, H., Stieglitz, J., Cummings, D. K., Wood, B., Emery Thompson, M., Raichlen, D., Beheim, B., Yetish, G., Kaplan, H., & Gurven, M. (2023). Energetic costs of testosterone in two subsistence populations. *American Journal of Human Biology*, *35*(11), e23949. https://doi.org/10.1002/ajhb.23949

van der Walt, L. A., Wilmsen, E. N., Levin, J., & Jenkins, T. (1977). Endocrine studies on the San ('Bushmen') of Botswana. *SA Medical Journal*, *52*, 230–232.

Vanjaarsveld, A. S., & Skinner, J. D. (1991). PLASMA ANDROGENS IN SPOTTED HYAENAS (CROCUTA-CROCUTA)—INFLUENCE OF SOCIAL AND REPRODUCTIVE DEVELOPMENT. *Journal of Reproduction and Fertility*, *93*(1), 195–201. (WOS:A1991GG00600023).

Vierhapper, H., Nowotny, P., & Waldhausl, W. (1997). Determination of testosterone production rates in men and women using stable isotope/dilution and mass spectrometry. *JOURNAL OF CLINICAL ENDOCRINOLOGY & METABOLISM*, *82*(5), 1492–1496. (WOS:A1997WX55100032). https://doi.org/10.1210/jc.82.5.1492

Von Engelhard, N., Kappeler, P. M., & Heistermann, M. (2000). Androgen levels and female social dominance in *Lemur catta*. *Proceedings of the Royal Society of London. Series B: Biological Sciences*, *267*(1452), 1533–1539. https://doi.org/10.1098/rspb.2000.1175

Wallace, K. M. E., Hart, D. W., Hagenah, N., Ganswindt, A., & Bennett, Nigel. C. (2023). A comprehensive profile of reproductive hormones in eusocial Damaraland mole-rats (Fukomys damarensis). *General and Comparative Endocrinology*, *333*, 114194. https://doi.org/10.1016/j.ygcen.2022.114194

Wang, L., Chen, G., Hou, J., Wei, D., Liu, P., Nie, L., Fan, K., Wang, J., Xu, Q., Song, Y., Wang, M., Huo, W., Jing, T., Li, W., Guo, Y., Wang, C., & Mao, Z. (2022). Ambient ozone exposure combined with residential greenness in relation to serum sex hormone levels in Chinese rural adults. *ENVIRONMENTAL RESEARCH*, *210*. (WOS:000777212100003). https://doi.org/10.1016/j.envres.2022.112845

Zihlman, A. L., & Bolter, D. R. (2015). Body composition in *Pan paniscus* compared with *Homo sapiens* has implications for changes during human evolution. *Proceedings of the National Academy of Sciences*, *112*(24), 7466–7471. https://doi.org/10.1073/pnas.1505071112

Zohdy, S., Gerber, B. D., Tecot, S., Blanco, M. B., Winchester, J. M., Wright, P. C., & Jernvall, J. (2014). Teeth, Sex, and Testosterone: Aging in the World's Smallest Primate. *PLOS ONE*, *9*(10), e109528. https://doi.org/10.1371/journal.pone.0109528
